# Caffeic Acid Phenethyl Ester (CAPE) Inhibits Growth of Chromosomally instable *bub1*Δ mutant in *Saccharomyces cerevisiae*

**DOI:** 10.1101/179994

**Authors:** Zeynep N. Azman, Aysel Kiyici, Mufide Oncel, H. Ramazan Yilmaz, Esra Gunduz, Mehmet Gunduz, Munira A. Basrai, Sultan Ciftci-Yilmaz

## Abstract

Chromosomal instability (CIN) is a hallmark of cancer cells. Spindle Assembly Checkpoint (SAC) proteins such as Bub1 monitor errors in chromosome segregation and cause cell cycle delay to prevent CIN. Altered expression of *BUB1* is observed in several tumor samples and cancer cell lines which display CIN. Caffeic Acid Phenethyl Ester (CAPE) which is an active component of propolis obtained from bee hives has anti-inflammatory antioxidant and anticarcinogenic properties. We used budding yeast *S. cerevisiae* as a model organism to investigate the molecular mechanism by which CAPE can inhibit the growth of cells with high levels of CIN. Here we show that CAPE leads to growth inhibition of *bub1*Δ strains. CAPE treatment suppressed chromosome mis-segregation in *bub1Δ* strain possibly due to apoptosis of chromosomally instable *bub1*Δ cells. We propose that CAPE may serve as a potential therapeutic agent for treatment of *BUB1* deficient cancers and other cancers that exhibit CIN.

## 1. Introduction

Chromosome instability (CIN) observed in >90% in solid tumors is one of the hallmarks of cancer cells (Rao et al. 2009). Several studies suggested that CIN is related with advanced stage tumors and resistance to chemotherapy (Thompson and Compton 2011). Exploring the potential of anticancer agents specifically targeting CIN in cancer cells offers a great potential for treatment of cancers. Caffeic acid phenethyl ester (CAPE), an active component of propolis obtained from bee hives has been reported to have anti-inflammatory, antioxidant, immunomodulatory, and anticarcinogenic properties (Borrelli et al. 2002; Cakir et al. 2011; Fadillioglu et al. 2010; Son and Lewis 2002; Yilmaz et al. 2004; Yilmaz et al. 2005; Iraz et al. 2006; Ozyurt et al. 2007; Koltuksuz et al. 2001; Michaluart et al. 1999). CAPE has also been shown to effectively inhibit cisplatin induced chromosome instability in rats (Yilmaz et al. 2010). However, the molecular mechanisms for growth inhibition by CAPE have not been well defined.

Bub1 (budding uninhibited by benzimidazole) is a component of Spindle Assembly Checkpoint (SAC) which is evolutionarily conserved from human to yeast (Kitagawa et al. 2001). SAC ensures faithful chromosome segregation by not allowing cells to undergo mitosis without correct kinetochore-microtubule attachment. Failure of SAC due to altered expression or deletion of *BUB1* results in increased CIN (Yuen et al. 2007) and several studies suggest that kinase defective *BUB1* plays a role in tumorigenesis (Kops et al. 2005). For example, four out of 19 colorectal cancer cell lines with CIN have mutations in *BUB1* (Cahill et al. 1998). Lymphoid leukemia and lymphoma cells also show deletions in the coding region of *BUB1* (Ru et al. 2002). Many tumors, especially the advanced stage tumors and cancer cell lines show altered expression of *BUB1* (Kops et al. 2005). Chromosome instability caused by reduced expression of *BUB1* results in thymic lymphoma in *p53^+/−^* mice and colonic tumors in *Apc^Min/+^* mice (Baker et al. 2009). Taken together, several studies provide direct evidence that a CIN phenotype due to mutations or altered expression of BUB1contributes to tumor formation in different cancers. Based on the role of *BUB1* in preventing CIN, we examined if *BUB1* can be exploited to examine the effect of new potential antitumor agents to target cells displaying a CIN phenotype.

We used budding yeast *Saccharomyces cerevisiae* as a model system to examine the effect of CAPE on chromosomally instable *bub1Δ* strain. Our results show that *bub1*Δ strains exhibit lethality and apoptosis when treated with CAPE. Furthermore, *bub1*Δ strains showed reduced chromosome segregation defects in the presence of CAPE. In summary our results provides mechanistic insights into the anticarcinogenic potential of CAPE and show that CAPE will be very effective in treatment of cancers that exhibit CIN.

## 2. Results and discussion

### 2.1. Growth inhibition of bub1Δ strain by CAPE

We used three different assays to investigate the effect of CAPE on growth of wild-type and *bub1Δ* cells. In the first assay, we performed growth assays of a fivefold serial dilution of cells on YPD with DMSO (control) or with 30 µg/ml CAPE in DMSO) incubated at 30°C. Wild-type cells did not show growth inhibition, however, *bub1Δ* cells showed very pronounced growth inhibition on medium containing CAPE (Figure 1A). The second assay quantified the growth inhibition phenotype by measurement of viability of wild-type and *bub1Δ* cells containing DMSO or 20 µg/ml CAPE in DMSO at 30°C. Percent survival was calculated by number of Colony Forming Units (CFU) on medium with and without CAPE. The *bub1*Δ strain showed significantly lower CFU when compared to wild-type strain on CAPE containing medium (Figure 1B). For the third assay, we measured the growth rate of wild-type and *bub1Δ* strains grown in liquid YPD with DMSO or 30 µg/ml CAPE in DMSO at 30°C was measured by OD_600_ every 3 hours. Treatment with CAPE reduced the growth rate of wild-type cells when compared to untreated cells. However, growth rate of the *bub1Δ* strain treated with CAPE was significantly decreased with no increase in OD_600_ after 12 and 24 hours compared to that for the wild-type strain grown in medium containing CAPE.

**Figure 1.**
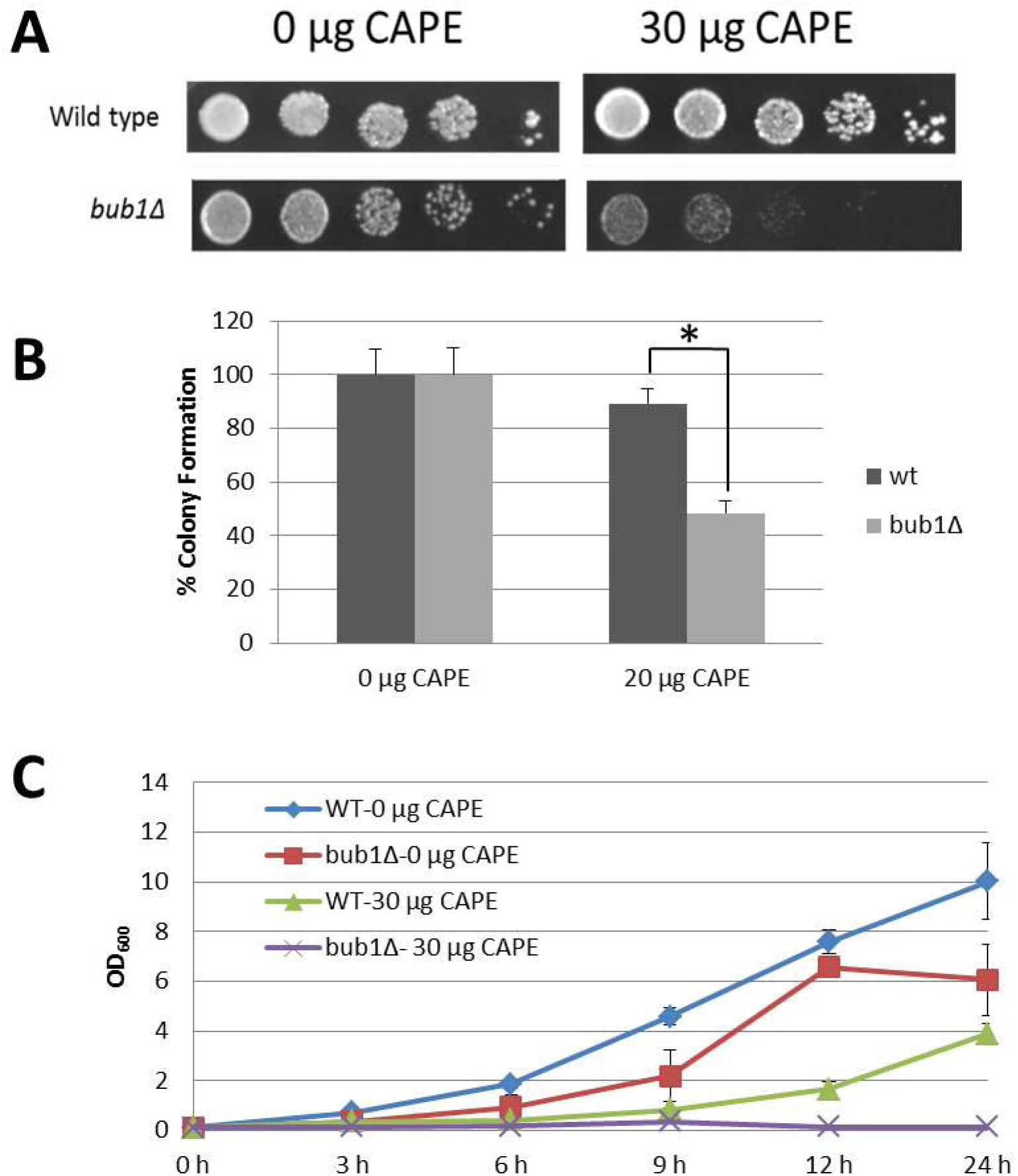
*bub1Δ* strain is sensitive to growth on CAPE containing medium. (a) Five-fold serial dilutions of wild-type and *bub1Δ* strains were spotted on YPD media containing either 0 or 30 µg/ml CAPE in DMSO. Plates were incubated at 30°C for two days and photographed. (b) Viability assays were done by measuring Colony Forming Units (CFU) of wild-type and *bub1Δ* strains plated on medium with DMSO or 20 µg/ml CAPE in DMSO at 30°C for 2-5 days (c) Growth rate of *bub1Δ* in liquid YPD media containing 0, 20 and 30 µg/ml CAPE in DMSO at 30°C was determined by optical density (OD_600_) measurement after every 3 hours. All experiments were done at least three times. Statistical significance was assayed with student’s t-test analysis, **p < 0.001, *p<0.05.

Cigut et al 2011 failed to observe any change in intracellular oxidation after treatment of yeast cells with CAPE (Cigut et al. 2011). Based on these result the authors concluded that lack of a cellular phenotype in their studies may be due accumulation of CAPE in membranes of yeast cells. Our results for growth inhibition of *bub1*Δ cells with CAPE treatment suggest that treatment with CAPE sensitizes cells predisposed to CIN.

### 2.2. Increased apoptosis due to chromosome fragmentation in CAPE treated bub1Δ cells

The growth inhibiton of *bub1*Δ strain to CAPE treatment prompted us to investigate if this is due to increased apoptosis. We used Terminal Deoxynucleotidyl Transferase Nick End Labeling (TUNEL) assay to investigate apoptotic effect of CAPE on wild-type and *bub1*Δ strains. Cells were grown until mid-logarithmic phase and treated with DMSO or 20µg/ml and 30µg/ml CAPE in DMSO for two hours. Cells were stained with 4',6-Diamidino-2-phenylindole (DAPI) to visualize the nucleus. *bub1*Δ strains showed higher incidence of chromosome fragmentation even without CAPE treatment when compared to wild-type cells. However, exposure to CAPE significantly increased chromosome fragmentation in the *bub1Δ* strain (Figure 2). Hence, we conclude that increased chromosome fragmentation may contribute to the lethality of *bub1Δ* cells in response to treatment with CAPE.

**Figure 2.**
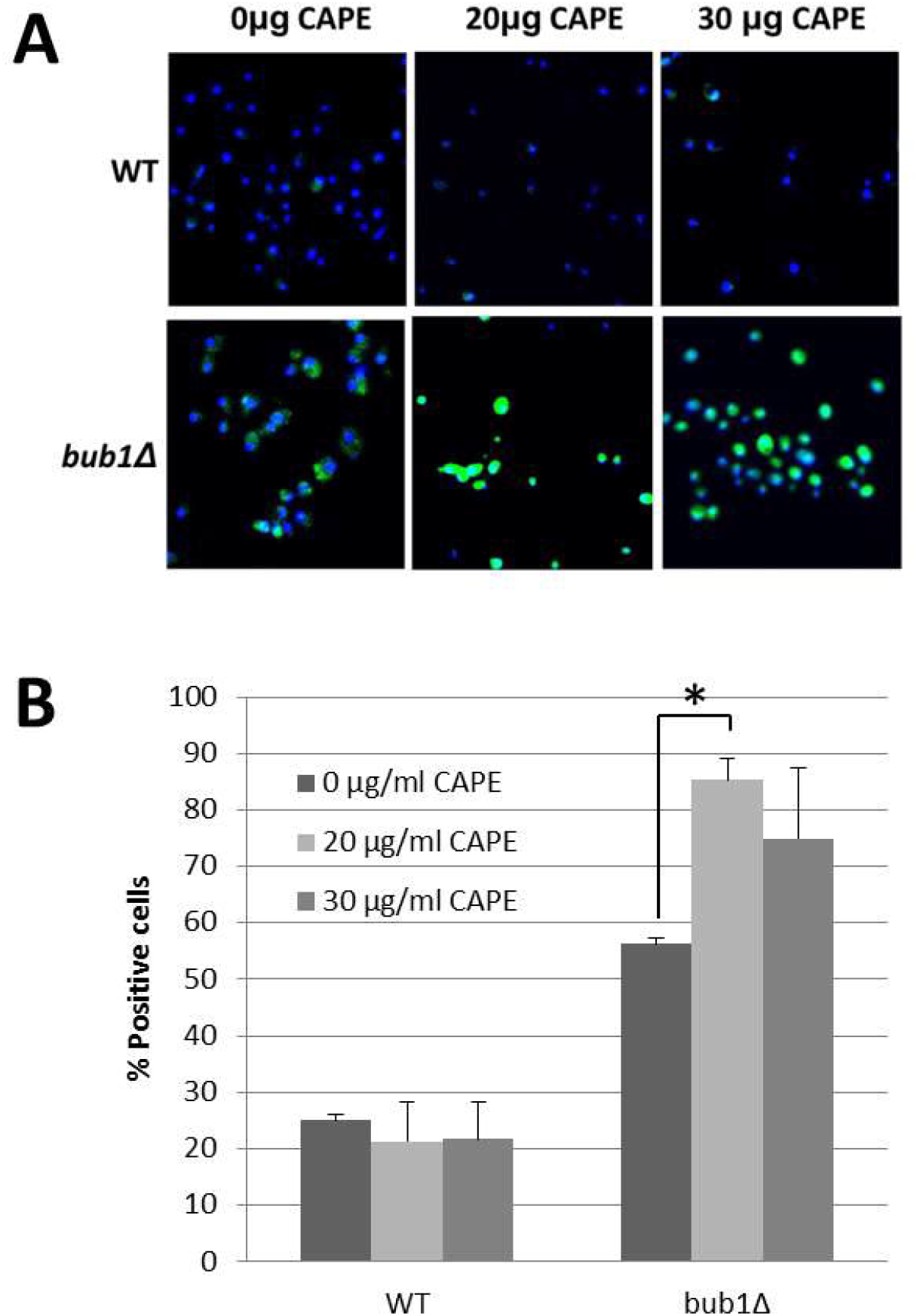
Increased apoptosis due to increased chromosome fragmentation of *bub1Δ* cells treated with CAPE. (a) Colocalization of Terminal Deoxynucleotidyl Transferase Nick End Labeling (TUNEL) and nucleus (DAPI) in wild-type and *bub1Δ* strains exposed to 0, 20 and 30 µg/ml CAPE (Blue: DAPI, Green: TUNEL). (b) Percent of TUNEL-positive cells exposed to 0, 20 and 30 μg/ml CAPE. Statistical significance was assayed with student’s t-test analysis, *p<0.05.

Our results provide mechanistic insights into the proapoptic effect of CAPE. Previous studies have shown that CAPE exhibits proapoptotic effect on tumors including C6 glioma (Lee et al. 2003), HCT116 human colorectal cancer (Wang et al. 2005), and MCF-7 breast cancer cell lines (Watabe et al. 2004). We propose that the increased CIN in these cancers makes them vulnerable to the effect of CAPE.

### 2.3. Decreased chromosome mis-segregation of bub1Δ strain after CAPE exposure

Previous studies have shown that CAPE treatment reduces cisplatin induced CIN in rat cells (Yilmaz et al. 2010). We used budding yeast to investigate if CAPE treatment reduces the CIN phenotype of *bub1*Δ strain. Chromosome segregation assays were done by measuring the segregation of a reporter chromosome with GFP at the centromere in wild-type and *bub1Δ* strain. Normal chromosome segregation results in the presence of one GFP-labeled chromosome in one nucleus, whereas defects in chromosome segregation results in more than one GFP-labeled chromosome in one nucleus. Treatment with CAPE (30µg/ml) resulted in about 50% reduction in chromosome segregation defects in *bub1Δ* strains but not a significant effect on CIN in wild-type strain (Figure 3). The reduced CIN of *bub1*Δ strain treated with CAPE are consistent with similar observations for reduced CIN in cisplatin treated rat cells (Yilmaz et al. 2010). It is possible that the *bub1*Δ cells with very high levels of CIN are eliminated by CAPE treatment and those that survive display lower levels of CIN. Future studies will allow us to decipher the molecular basis for the reduced CIN in CAPE treated *bub1*Δ cells.

**Figure 3.**
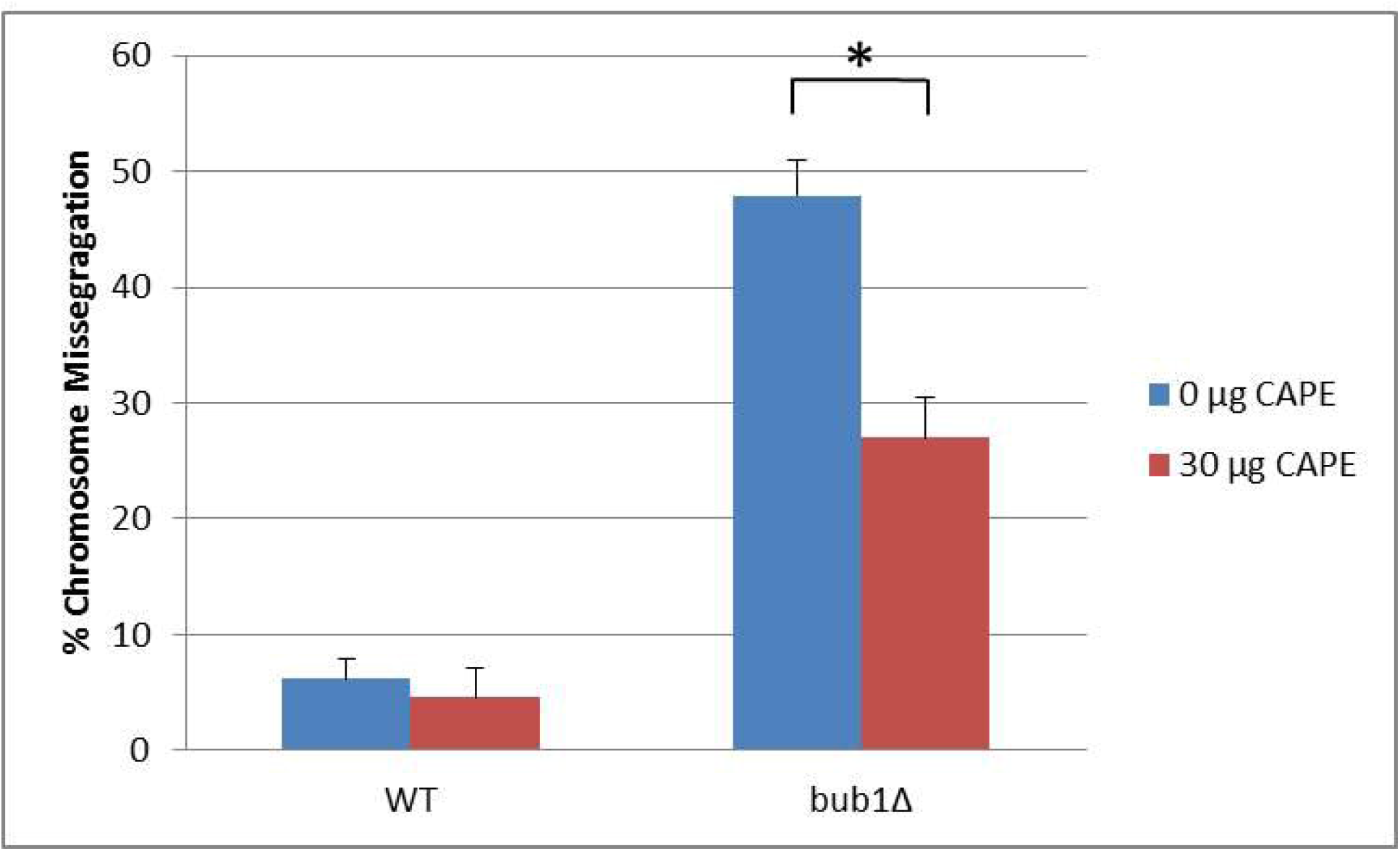
CAPE suppresses chromosome segregation defects in *bub1Δ* strain. Segregation of a GFP-labeled chromosome in wild-type and *bub1Δ* strains exposed to 0 and 30 μg/ml CAPE for four hours. Normal chromosome segregation is scored by the presence of GFP labeled chromosome within the nucleus of unbudded cells or in each of separated nuclei in large-budded cells. Defects in chromosome segregation is scored by the presence of more than one GFP labeled chromosome within a single nucleus or large budded cells in which one nuclear mass was devoid of GFP foci. Statistical significance was assayed with student’s t-test analysis, *p<0.05.

## 3. Experimental

### 3.1 Strains, media and culture

The following *Saccharomyces cerevisiae* strains obtained from Yeast Knockout Collection (Dharmacon, YSC1053) were used for growth and TUNEL assays: wild type-BY4741 (MAT a *his3Δ1 leu2Δ0met15Δ0 ura3Δ0), bub1*Δ (MATA *his3Δ1 leu2Δ0 met15Δ0 ura3Δ0 bub1::kanMX4)*, DDY1925 (MATA *his3-Δ200 ura3-52 ade2-1 HIS3::pCu-lac1-GFP leu2-3,112::lacO::LEU2)* (Cheeseman et al. 2002). The *bub1*Δ strain used for chromosome segregation assay was generated by replacing endogenous *BUB1* with the KanMX cassette in the DDY1925 using homologous recombination. The KanMX cassette was amplified from the BY4741 *bub1*Δ strain by the PCR method with the following primers: Bub1 Fwd ( 5’-TGAATGTTAACGCTGACCAGG-3’) and Bub1 Rvs (5’-ACCAAAAAGTCACCTATGCGG-3’). Gene replacement was confirmed by PCR with the following primers: Bub1 Fwd ( 5’-TGAATGTTAACGCTGACCAGG-3’) and KanB (5’-CTGCAGCGAGGAGCCGTAAT-3’). The following media used for cell growth unless otherwise is indicated: YPD (1% yeast extract, 2% bactopeptone, and 2% glucose), solid YPD (1% yeast extract, 2% bactopeptone, 2% glucose, and 2% agar). *bub1Δ* transformants for the chromosome segregation assay were selected on YPD medium with 200 µg/ml G418 (Sigma, A1720). CAPE used in all experiments was obtained from Sigma (C8221) and dissolved in Dimethyl sulfoxide (DMSO).

### 3.2. Growth assays

For growth assays, three different methods were utilized. For all assays logarithmic phase cells were used. For the first assay, a five-fold serial dilution of cells was spotted on solid YPD with DMSO or 30 µg/ml CAPE in DMSO and incubated at 30°C for two days. For measurement of colony forming units (CFU) cells were spread on YPD with DMSO or YPD with 20 µg/ml CAPE in DMSO and incubated at 30°C for two-five days. For growth rate analysis, the cells were grown in YPD with DMSO or 30 µg/ml CAPE in DMSO at 30°C and the optical density (OD_600_) was measured in every 3 hours.

### 3.3. Chromosome segregation assay

Chromosome segregation assay was performed as described in Boeckmann et al., 2013 with some modifications (Boeckmann et al. 2013). DDY1925 and *bub1Δ* strains were grown to logarithmic phase in YPD medium containing 0.8 mM adenine to reduce background fluorescence and 250 µM CuSO_4_ to induce expression of the LacI-GFP fusion reporter at 30°C. Cells were incubated with DMSO or 30µg/ml CAPE in DMSO for four hours at 30°C and fixed for 10 minutes in 4% formaldehyde at room temperature. After two washes with phosphate-buffered saline (PBS), the cells were resuspended in PBS containing 10 µg/ml DAPI (4',6-diamidino-2-phenylindole). The cells were visualized in a fluorescent microscope. Both single cells and large budded cells with clear nuclear separation, identified by DAPI fluorescence, were used for scoring chromosome missegregation. One GFP-labeled chromosome in one nucleus scored as normal chromosome segregation, whereas more than one GFP-labeled chromosome in one nucleus scored as chromosome mis-segregation.

### 3.4. TUNEL assay

DNA strand breaks as indicator of apoptosis was investigated by Apop Tag Fluorescein In Situ Apoptosis Detection Kit (Millipore, S7110). Yeast cells were grown until mid-logarithmic phase and incubated with 0, 20 or 30 µg/ml CAPE in DMSO. Cells were washed with water and fixed with 3.7% (vol/vol) formaldehyde in 0.1M phosphate citrate buffer (0.1 M dibasic sodium phosphate, 0.05 M sodium citrate, pH 5.8) for 30 minutes at room temperature. For cell wall digestion, cells were washed two times with 0.1 phosphate citrate buffer and incubated with 0,5U/µl Lyticase from *Arthrobacter luteus* (Alfa Aesar, J63195) at 37°C until spheroplast formation was observed in the majority of the cells. Cell suspension was applied to a microscope slide and allowed to dry at room temperature. The slides were rinsed with PBS and incubated with ethanol: acetic acid (2:1) solution at -20°C for 5 minutes. After rinsing the slides with PBS, the samples were incubated with the following buffers consecutively; equilibration buffer for 10 seconds, TdT enzyme at 37°C for 1 hour, stop/wash buffer for 10 minutes and DAPI. The slides were mounted and visualized in a fluorescent microscope.

### 3.5. Statistical Analysis

In all experiments, statistical significance was assayed with student’s t-test analysis.

## 4. Conclusion

In summary, we provide the first evidence for correlation between increased efficacy of growth inhibition by CAPE treatment and cells with a CIN phenotype. Our results show increased growth inhibition of CAPE treated *bub1*Δ strain which displays CIN. CAPE treatment significantly inhibits growth and this contributes to reduced viability of the *bub1Δ* cells compared to wild-type cells. Intriguingly CAPE treatment suppressed chromosome segregation defects in surviving *bub1Δ* cells. Given the evolutionarily conservation of pathways of chromosome segregation our results with budding yeast provide an opportunity to further investigate the potential of CAPE as a potential chemotherapeutic agent for inhibition of cancers with a CIN phenotype such as those with defects in *BUB1*.

## Acknowledgement

This work was supported by the Scientific and Technological Research Council of Turkey (TUBITAK) (112S254).

## Conflict of Interest

The authors declare no conflict of interest.

